# Are narrow-ranging species doomed to extinction? Projected dramatic decline in future climate suitability of two highly threatened species

**DOI:** 10.1101/2021.03.30.437650

**Authors:** Nicolas Dubos, Frederique Montfort, Clovis Grinand, Marie Nourtier, Gregory Deso, Jean-Michel Probst, Julie Hanta Razafimanahaka, Raphali Rodlis Andriantsimanarilafy, Eddie Fanantenana Rakotondrasoa, Pierre Razafindraibe, Richard Jenkins, Angelica Crottini

## Abstract

Narrow-ranging species are usually omitted from Species distribution models (SDMs) due to statistical constraints, while they are predicted to be particularly vulnerable to climate change. The recently available high-resolution environmental predictors, along with recently developed methods enable to increase the eligibility of narrow-ranging species for SDMs, provided their distribution is well known. We fill a gap of knowledge on the effect of predicted climate change on narrow-ranging species. We modelled the distribution of the golden mantella frog *Mantella aurantiaca* and the Manapany day gecko *Phelsuma inexpectata*, for which the distribution of their occurrence records is well documented. Our modelling scheme included a range of processes susceptible to address statistical issues related to narrow-ranging species. We predict an alarming decline in climate suitability in the whole current distribution area of both species by 2070, potentially leading to a complete extinction in most scenarios. We identified the areas with the best climate suitability in the future, but these remain largely suboptimal regarding species climatic niche. The high level of habitat fragmentation suggests that both species likely need to be at least partly translocated. Climate change may not only drive range contractions or distribution shifts in narrow-ranging species, but may lead to the complete extirpation of suitable environments across their entire region. This study suggests that the level of threats of narrow-ranging species already identified as threatened may be underestimated, especially in heterogeneous tropical areas. We stress the need to develop sampling campaigns and implement proactive actions for narrow-ranging species in the tropics.

## Introduction

Climate change is predicted to become the main driver of biodiversity loss it in the next decades (Bellard *et al*., 2012). Species Distribution Models (SDMs) are probably the most common approach used to predict the impact of future climate change on species. They are used to predict current and future environmental suitability, and provide guidelines for the identification of priority areas for protection (Leroy *et al*., 2014), habitat restoration and species (re)introduction/translocations (e.g., Bellis et al., 2020; Draper et al., 2019; Butt et al., 2020; Westwood et al., 2020). Habitat restoration and translocations are two ecological engineering techniques enabling the restoration of depleted populations. Translocation programs will become increasingly needed in the face of climate change, especially for species with small distribution ranges (Thomas, 2011). In this regard, SDMs help identifying suitable receptor sites that meet the species’ habitat requirements while accounting for climate suitability (Bellis et al., 2020). In highly degraded environments a combination of habitat restoration and translocation may be needed to avoid species extinctions. This may be the case in highly fragmented tropical systems, where the number of narrow-ranging species is higher and where climate change effects are expected to be stronger (Tewksbury et al., 2008).

The impact of future climate change is largely understudied in endangered narrow-ranging species. This is mainly due to the difficulty to model their current distribution and project their future because of low sample sizes and subsequent little spatial replicates when fitted on climate data (Botts *et al*., 2013; Platts et al., 2014; Breiner et al., 2015; Galante et al., 2018). Both these factors lead to statistical constraints that withdraw these species from eligibility for SDMs. However, the omission of narrow-ranging species in SDMs may be problematic in terms of conservation planning, because the area encompassing their distribution may be downplayed (Platts *et al*., 2014). Although narrow-ranging species are known to be more vulnerable to climate change (Pearson *et al*., 2014), few studies have provided quantitative assessment of climate change impacts on these (but see Alamgir *et al*., 2015; Zhang *et al*., 2020).

The recent availability of high-resolution climatic data (e.g., Fick & Hijmans, 2017; Karger et al., 2017), along with high-resolution land cover data (e.g., Vieilledent et al., 2018) is offering new opportunities for modelling the distribution of these species (Lannuzel et al., 2021). However, despite a probable increase in statistical power, there is still the possibility for SDMs to produce misleading results due to spatial sampling bias (Phillips *et al*., 2009). The effect of sample bias may be particularly strong in rare (or poorly known) species with small sample sizes, because models are more influenced by each occurrence data that is used (Pearson *et al*., 2007). A number of techniques were recently developed for data-poor species. For instance, jackknife procedures (i.e. leave-one-out; Pearson et al. 2007) improves model assessments and Ensemble of Small Models (ESMs) enable to deal with model complexity while keeping sufficient explanatory power (Breiner et al. 2015). Other techniques are dedicated to sampling bias correction, often implying the filtering of occurrence or environmental data (Gábor *et al*., 2019) or non-random pseudo-absence selection (Phillips *et al*., 2009). However, data filtering can become problematic for species with low sample sizes (Vollering et al., 2019), especially when species distribution is highly localised (Inman et al., 2021) and is not recommended in absence of evidence of bias in occurrence data (Gábor et al., 2019). Similarly, non-random pseudo-absence selection is not always effective (Dubos et al., 2021) and tends to make predictions worse in narrow-niche species (Inman et al., 2021). However, the restriction of the background area (where pseudoabsences are generated), i.e. background thickening, can be advantageous for sample bias correction when data is scarce without discarding presence records (Acevedo et al. 2012; Vollering et al. 2019). On the other hand, the reliability of an SDM is more driven by the quality of the data than the implementation of models (Araùjo et al., 2019). Therefore, the best option for narrow-ranging SDMs may be to (1) use all the available methodological tools to deal with statistical issues, and (2) select species for which the distribution is well known.

Given the predicted magnitude of climate change, along with the narrow thermal tolerance of tropical species (Tewksbury et al., 2008), we may not only expect a reduction or a geographical shift in suitable conditions for narrow-ranging species, but the extirpation of suitable conditions across the entirety of their distribution range. Here we fill a gap of knowledge regarding the impact of future climate change on narrow-ranging species using two species for which the distribution is particularly well documented. These were the Manapany day gecko *Phelsuma inexpectata*, classified as Critically Endangered (IUCN & MNHN, 2010) and the golden mantella frog *Mantella aurantiaca* formerly classified as Critically Endangered (Vences & Raxworthy, 2008), now classified as Endangered after the inclusion of one locality record which increased its extent of occurrence (IUCN SSC Amphibian Specialist Group 2020). Both species are in continued decline (Probst & Turpin, 1997; Crottini et al., 2019), live in highly fragmented areas (respectively in Reunion Island and central Madagascar) and are in urgent need for conservation actions. Given the important, long-term efforts invested to document their distribution, we assume that the geographic information for these species is nearly comprehensive and unbiased. We tested whether climate change will ‘only’ drive range reductions/shifts, or lead to a total extirpation of their suitable areas. We eventually identify the most suitable candidate areas for habitat restoration and translocation across their respective regions.

## Methods

### Occurrence data

#### Phelsuma inexpectata

The Manapany day gecko is endemic to the south of Reunion Island. We retrieved 31 occurrence data from literature (Bour *et al*., 1995). Since then, the surroundings of the known distribution range of the species were regularly visited (an 11 km-long coastal band; Fig. 1). Two localities corresponding to introduced populations were identified west of the current range, which we added to the data (Deso, 2001; Porcel *et al*., 2021). Recent sampling campaigns enabled to find additional occupied habitats (Dubos 2010) but did not add any occurrence point after aggregation at the resolution of the environmental variables (30 arc seconds). Therefore, we assume that the sample occurrence of the species is nearly comprehensive. The total number of 30 sec. occupied pixels resulted in 15 presence points.

**Fig. 1.**
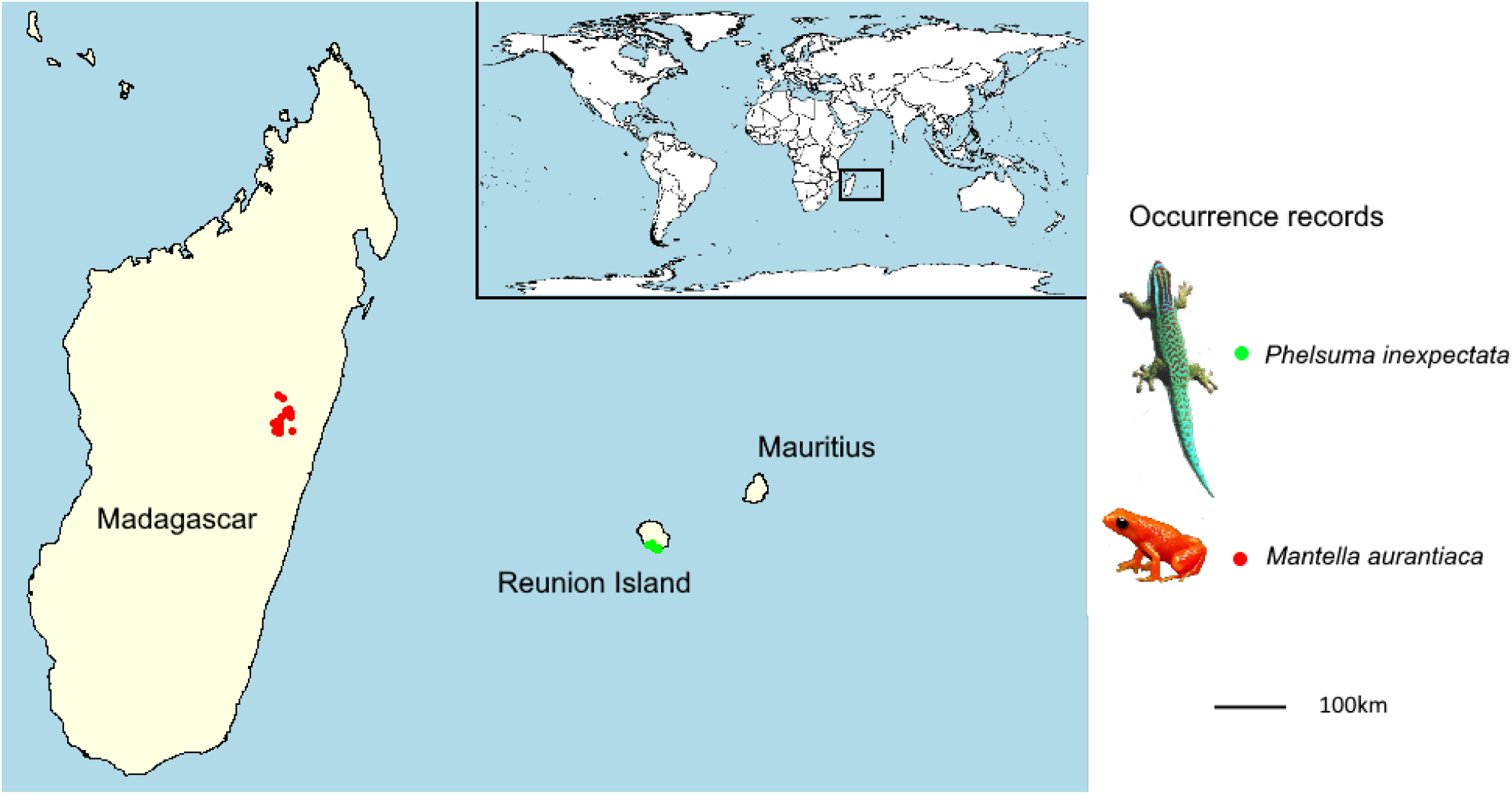
Study area and species, with occurrence points.

#### Mantella aurantiaca

The golden mantella frog is distributed in central-Eastern Madagascar (region of Moramanga; Fig. 1). We obtained 131 occurrence data from Piludu *et al*. (2015). Those included compiled published data from surveys conducted between 2001 and 2007 (Bora *et al*., 2008; Randrianavelona *et al*., 2010) and new locations from additional surveys conducted between 2008 and 2013. More recent surveys conducted between 2014 and 2019 enabled to add 39 occurrences. The region has been extensively surveyed and it is very likely that most occupied habitats were identified. After aggregation to match the resolution of environmental data, sample size resulted in 101 occurrence points.

### Climate data

We used the 19 bioclimatic variables for 30 arc sec (approximately 900m) resolution of the current climate data and of the 2070 projections from CHELSA (Karger et al., 2017; Fig. S1, S2). We decided to include all the 19 variables because both temperature and precipitation are related to the species’ biology, including those related to indices of variability (e.g., cyclones drive mortality in *Phelsuma* and heavy rains drive reproduction in *Mantella*; Vinson, 1975; Randrianavelona et al., 2010). We used three Global Circulation Models (GCMs; i.e., BCC-CSM1-1, MIROC5 and HadGEM2-AO) and two greenhouse gas emission scenarios (the most optimistic RCP26 and the most pessimistic RCP85; Fig. S3, S4). We also ran sets of models using the WorldClim Global Climate Data (Fick & Hijmans, 2017), with the same GCMs and scenarios to account for potential effect of the baseline data in our predictions.

### Distribution modelling

We modelled and projected species distributions with the biomod2 R package (Thuiller et al. 2009), using 10 modelling techniques: generalised linear model (GLM), generalised additive model (GAM), classification tree analysis (CTA), artificial neural network (ANN), surface range envelop (SRE, as known as BIOCLIM), flexible discriminant analysis (FDA) and random forest (RF), Multiple Adaptive Regression Splines (MARS), Generalised Boosting Model (GBM) and Maximum Entropy (MaxEnt). For machine learning modelling techniques, hyperparameters were set to their default values implemented in biomod2 (see https://www.rdocumentation.org/packages/biomod2/versions/3.5.1/topics/BIOMOD_ModelingOptions). We generated five different sets of randomly-selected pseudo-absences (random generation is recommended for rare and specialised species; Inman et al., 2021). We selected one variable per group of inter-correlated variables to avoid collinearity (Pearson’s r > 0.7) and assessed the relative importance of each variable kept with 10 permutations per model replicate (total = 500). The variables included in the final models were those with a relative importance > 0.2 across at least 50% of model runs (these varied with the baseline climate and the background extent; Table 1). We show the fitted predicted values for each individual model, and a mean smoothed response curve. We predicted species distributions with an ensemble of small models approach (ESM; Breiner et al., 2015). We ran sets of bivariate models, i.e. including all pairwise combinations of the selected variables, and produced an ensemble model with the mean predictions across all models weighted by their respective AUC (see below). This method is advocated for rare species and enables to reduce model complexity without reducing the explanatory power. We set three runs of cross validation (except for the jackknife procedure; see below). We ran a first set of models (1), hereafter referred to as ‘Base’ setting, with 1000 pseudo-absences selected from a background covering the entire respective island of each species (i.e. all island cells). We ran a second set (2) where the 1000 pseudo-absences were down-weighted to equal presence data (setting prevalence to 0.5) referred to as ‘equal total weight’ (ETW; Liu et al. 2019). We reperformed the first two model sets but with a background covering the southern part of Reunion Island and the eastern forested part of Madagascar (see ‘Land use data’ section), without, and with pseudo-absence down-weighting, respectively referred to as (3) ‘Restricted background’ and (4) ‘ETW-Restricted background’. Eventually, we ran a (5) Base model set with the Worldclim baseline climate instead of Chelsa, referred to as ‘Worldclim baseline’. For both species we present five ensemble models of current distribution (of varying background, prevalence and baseline climate), and 30 ensemble models of projected future distribution (of varying background, prevalence, baseline climate, GCM and RCP). We did not account for species dispersal ability, because the purpose was not to predict potential shifts, but to assess how suitable will be the climate in the future in order to identify candidate sites for restoration and translocation.

#### Model evaluations

Ideally, performance evaluations are based on block-cross validation to limit spatial autocorrelation at large scales. In our case, species distributions are highly localised and the use of spatial splits would result in strong unbalances between blocks. We therefore used a random partitioning for *M. aurantiaca* and a jackknife for *P. inexpectata*. The jackknife procedure is appropriate for small sample sizes (Pearson *et al*., 2007; Galante *et al*., 2018). This approach consists in running n iterations corresponding to the number of occurrence data, removing one occurrence for each run of calibration. Models are evaluated with the withheld occurrence. We did not used it for *M. aurantiaca* given that the number of occurrence data was sufficient for a standard procedure, and used an 80% calibration subset (random partitioning), with five cross-validation runs. We assessed model performance using the Area Under the Operative Curve (AUC), the True Skill Statistics (TSS) and the Boyce index (the latter was not computed for the jackknife procedure, because this index is based on Spearman’s coefficients and cannot be estimated on the basis of a single location). Poorly performing models could result from calibration subsets falling into different conditions than evaluation subsets. For ensemble models (mean predictions), we excluded models for which the AUC was below 0.9 and for which the Boyce index was below 0.5 (Gillard et al., 2017).

We performed a multivariate environmental similarity surfaces analysis (MESS) to determine whether models are well informed for projections on novel (future) data. We eventually quantified the uncertainty related to model input parameters (modelling technique, pseudo-absence distribution, number of pseudoabsences generated, pseudo-absence down-weighting, GCMs, RCPs and baseline climate data) by computing the standard deviation of the suitability scores between model predictions.

### Land use data

We accounted for the habitat requirements of our model species by applying a filter to the projected climate suitability based on land use and land cover data (e.g., Gillard *et al*., 2017). This enabled to minimise model complexity while remaining biologically realistic and relevant for conservation applications.

Reunion Island – We used very high-resolution land cover categories (Urban, agricultural, natural, water) at 1.5m resolution (resampled at 15m for computing purposes) derived from remote sensing (Dupuy and Gaetano, 2019; Fig. S5). Since *P inexpectata* can be found in both urbanised and natural areas and avoids agricultural areas (Probst & Turpin, 1997), we excluded the latter land use type only. This is relevant also for conservation applications because habitat restoration is more difficult to implement in agricultural areas.

Madagascar – *Mantella aurantiaca* is exclusively found in swampy forested areas (Randrianavelona *et al*., 2010). We filtered our suitability maps to include only cells corresponding to rainforests. We obtained forest cover data for two periods (1990 and 2017) from a study that combined global tree cover loss data with historical national forest (resolution 30m; Vieilledent et al., 2018; Fig. S6). We used forest cover of the year 1990 for current distribution models because it corresponds approximately to the period of the oldest occurrence data (some areas were deforested since then). For future projections, we used the latest forest cover data available, i.e. 2017.

## Results

### Variable selection and model performance

Three to four variables were selected in every cases (no change between prevalence settings; see details in Table 1; Fig. S7, S8). The current distribution of both species was well predicted (across five model sets: *P inexpectata*: mean AUC = 0.96, mean TSS = 0.91; *M. aurantiaca*: mean Boyce = 0.82, mean AUC = 0.97, mean TSS = 0.94; Fig. S9, S10). We excluded 5076 poorly performing models out of 18750 for *P inexpectata*, and 343 out of 4140 for *M. aurantiaca*. The median suitability score at the presence points for *P inexpectata* was 836 (911 when removing the two introduced populations) and 783 for *M. aurantiaca*. Models consistently identified the current distribution range between all runs for both species, as shown by the uncertainty map (SD intermodel suitability scores at the presence points of 168 for *P inexpectata* and 113 for *M. aurantiaca*; Fig. S11, S12).

**Table 1.** Selected variables for two narrow-ranging species in Reunion Island and Madagascar. Variables were obtained from CHELSA (Karger et al., 2017) and Worldclim (Fick & Hijmans, 2017), and selected on the basis of 10 permutations per modelling technique and pseudo-absence set (total = 500).

| Species | Model set | Selected variables |
| --- | --- | --- |
| *Phelsuma inexpectata* | Base | Bio1, Bio15, Bio18 |
|  | Restricted background | Bio1, Bio15, Bio18 |
|  | Worldclim baseline | Bio1, Bio3, Bio14, Bio16 |
| *Mantella aurantiaca* | Base | Bio1, Bio3, Bio4, Bio5 |
|  | Restricted background | Bio1, Bio18, Bio19 |
|  | Worldclim baseline | Bio10, Bio11, Bio14, Bio16 |

### Future climate suitability

All scenarios indicated an important decrease in climate suitability in the entirety of the current range of both species by 2070 (Fig. 2–7). On average across scenarios and GCMs, suitability scores decreased by 59% for *P inexpectata* (Fig. 8) and 73% for *M. aurantiaca* at the presence points (Fig. 9). For *M. aurantiaca*, only one scenario out of 30 showed high suitability values within the current range (i.e. the MIROC5 GCB, RCP85, Chelsa baseline, restricted background).

**Fig. 2.**
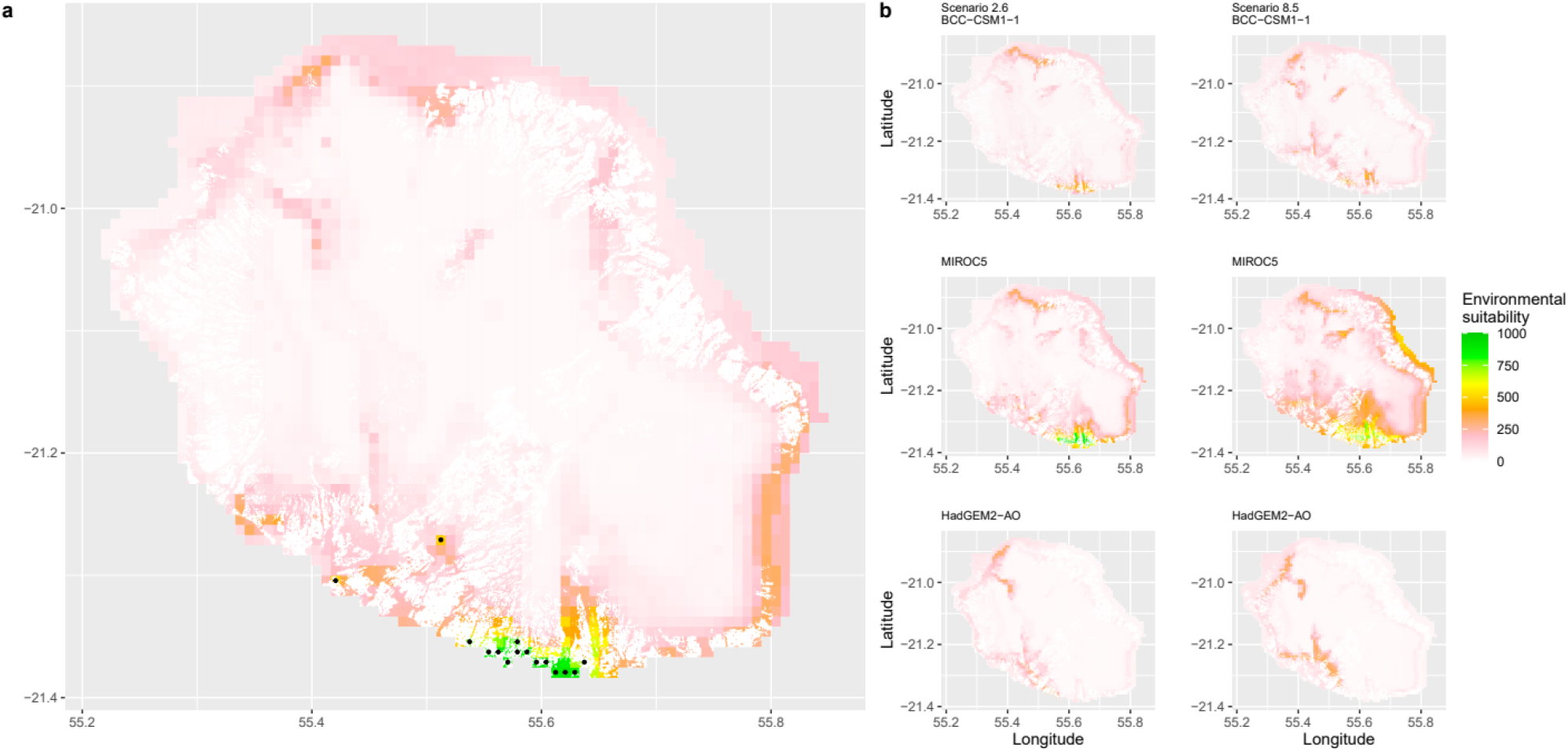
Current (a) and projected future (b) environmental suitability for *Phelsuma inexpectata* using Chelsa as baseline climate data, no prevalence setting and the whole island as a background. Predictions were filtered to remove agricultural areas. X and Y axes represent the coordinates (WGS84). Black points represent the occurrence data. Future projections (2070) were estimated from three Global Circulation models (GCM) and two RCP scenarios. Left panels represent the most optimistic scenario (RCP26) and right panels represent the most pessimistic scenario (RCP85). Models produced similar results when setting prevalence to 0.5 (Equal total weights).

**Fig. 3.**
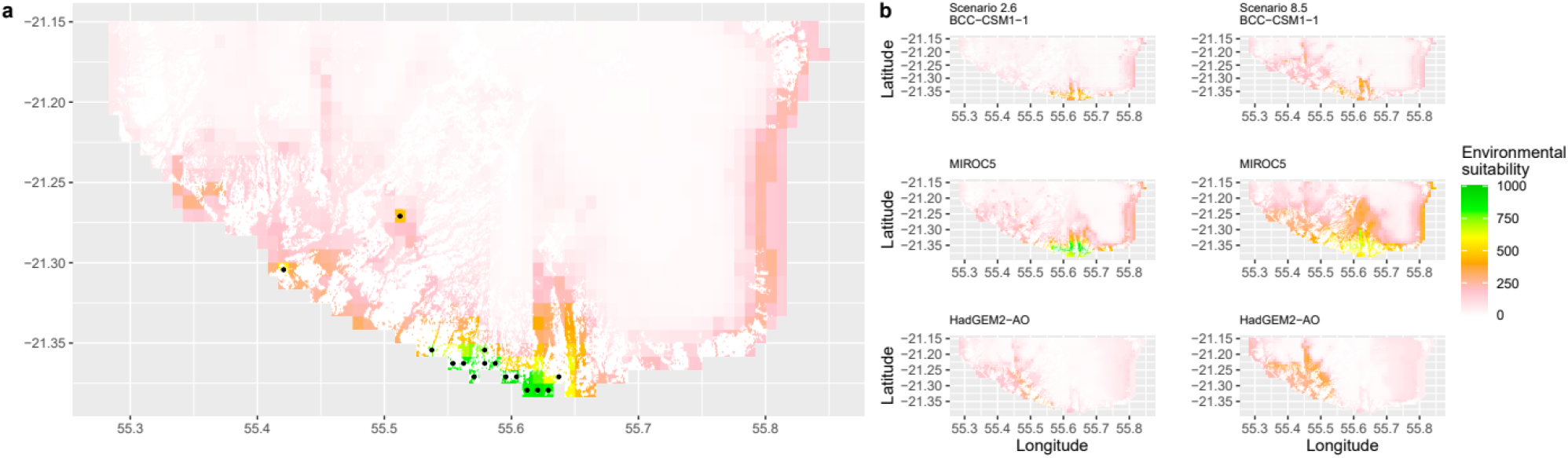
Current (a) and projected future (b) environmental suitability for *Phelsuma inexpectata* using Chelsa as baseline climate data, no prevalence setting and restricted background. Predictions were filtered to remove agricultural areas. X and Y axes represent the coordinates (WGS84). Black points represent the occurrence data. Future projections (2070) were estimated from three Global Circulation models (GCM) and two RCP scenarios. Left panels represent the most optimistic scenario (RCP26) and right panels represent the most pessimistic scenario (RCP85). Models produced similar results when setting prevalence to 0.5 (Equal total weights).

**Fig. 4.**
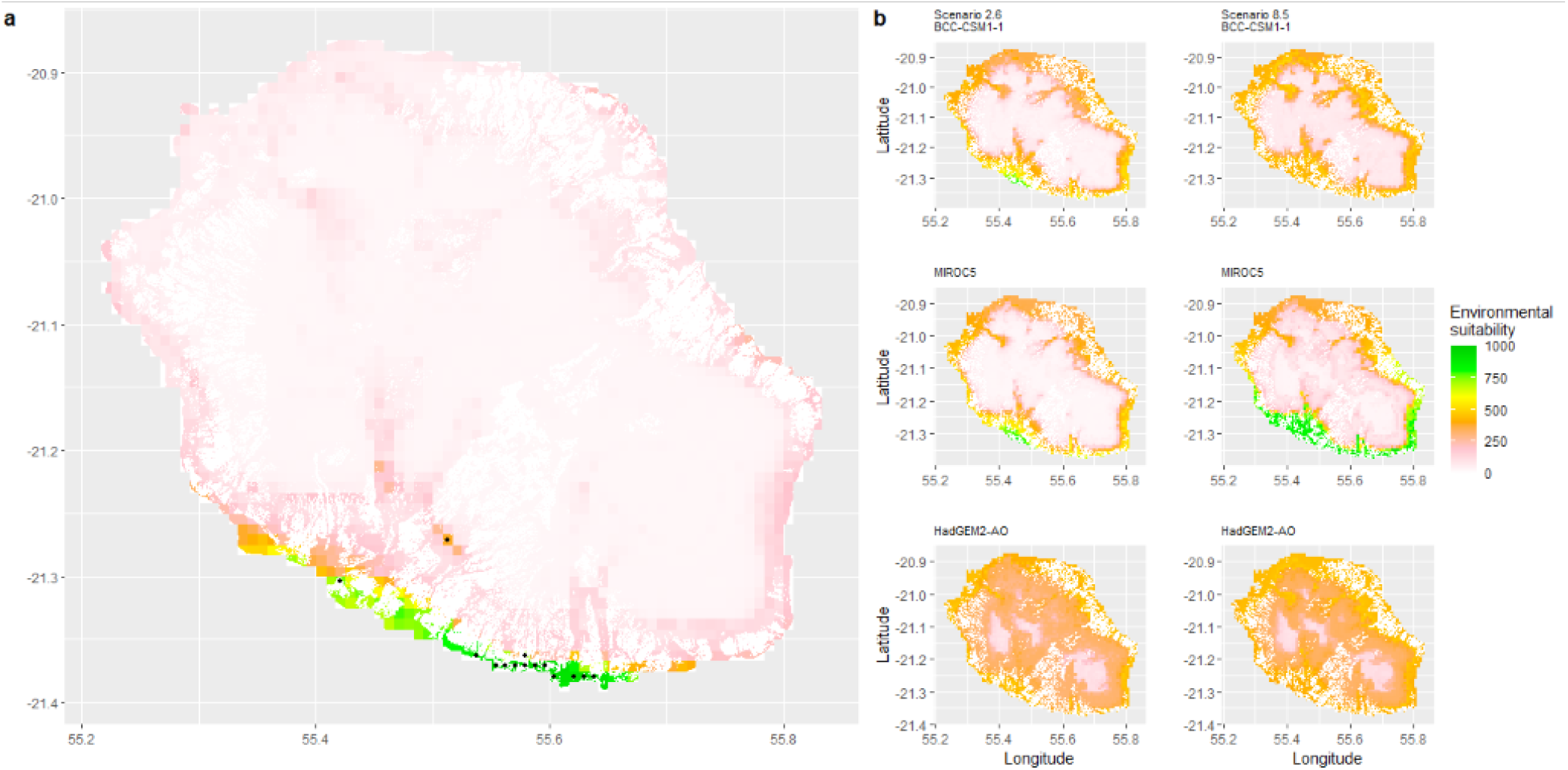
Current (a) and projected future (b) environmental suitability for *Phelsuma inexpectata* using Worldclim as baseline climate data. Predictions were filtered to remove agricultural areas. X and Y axes represent the coordinates (WGS84). Black points represent the occurrence data. Future projections (2070) were estimated from three Global Circulation models (GCM) and two RCP scenarios. Left panels represent the most optimistic scenario (RCP26) and right panels represent the most pessimistic scenario (RCP85).

**Fig. 5.**
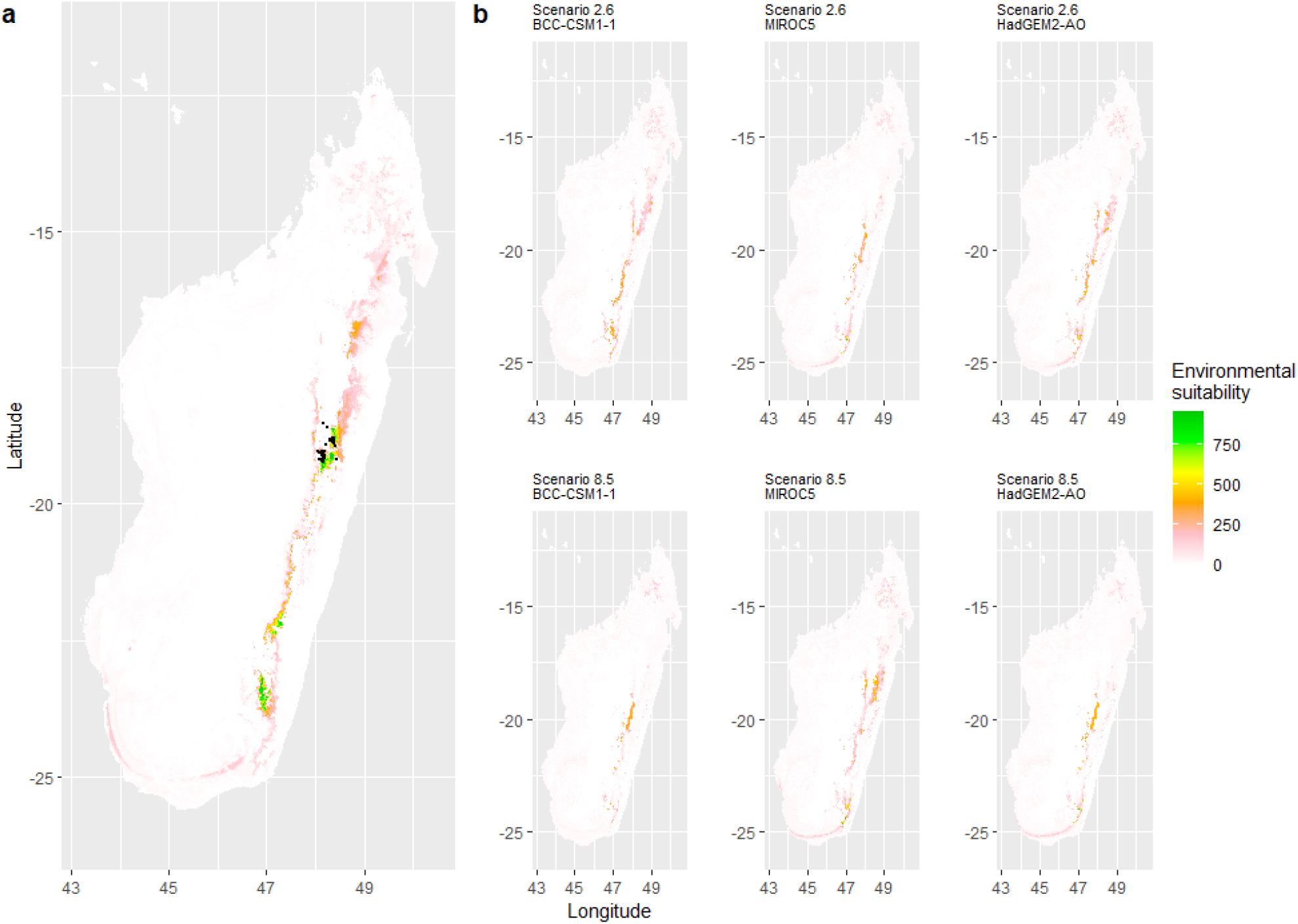
Current (a) and projected future (b) environmental suitability for *Mantella aurantiaca* using the Chelsa baseline climate, no prevalence setting and the whole island as a background. Predictions were filtered to include only forested areas. X and Y axes represent the coordinates (WGS84). Black points represent the occurrence data. Future projections (2070) were estimated from three Global Circulation models (GCM) and two RCP scenarios. Top panels represent the most optimistic scenario (RCP26) and bottom panels represent the most pessimistic scenario (RCP85). Models produced similar results when setting prevalence to 0.5 (Equal total weights).

**Fig. 6.**
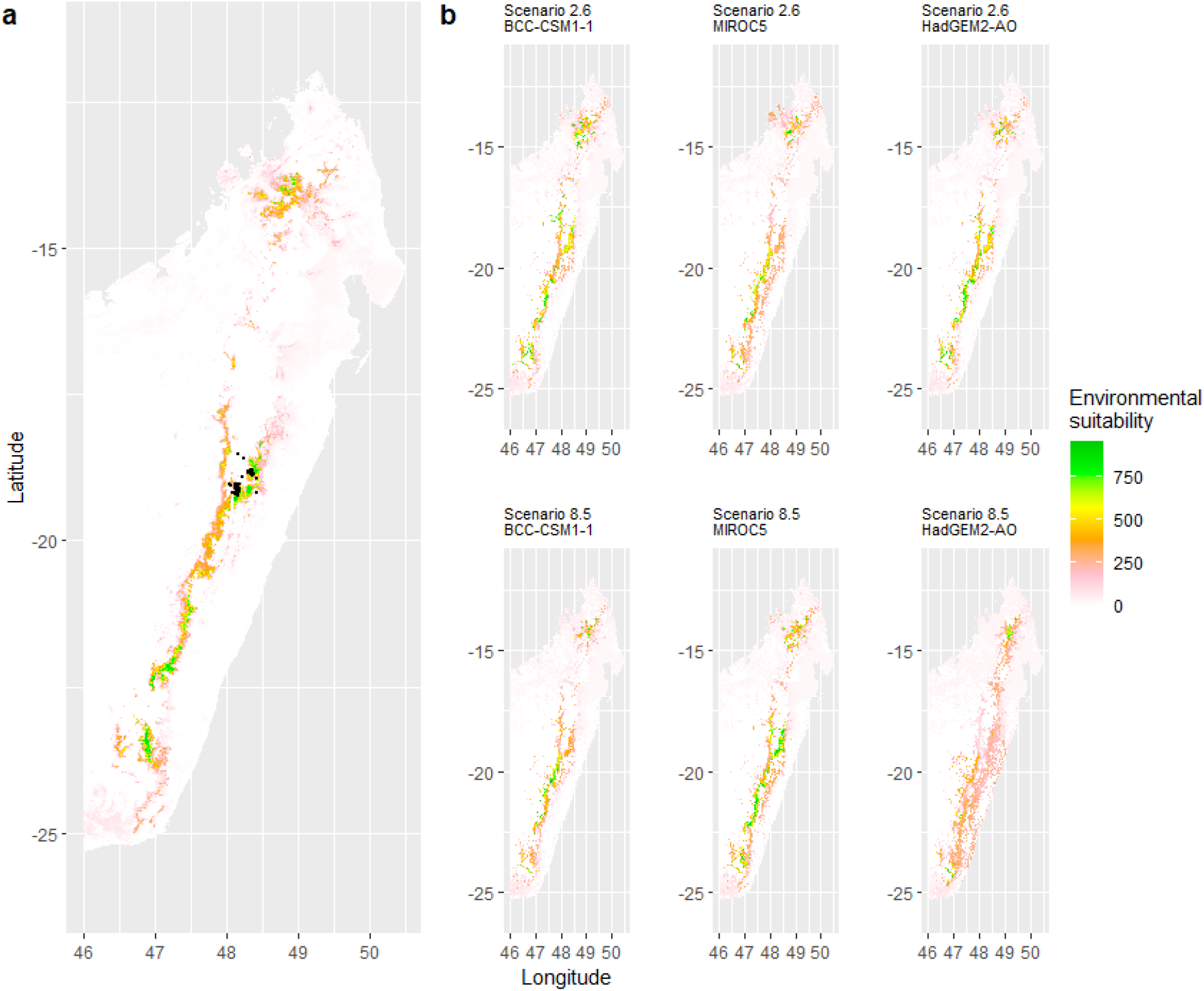
Current (a) and projected future (b) environmental suitability for *Mantella aurantiaca* using the Chelsa baseline climate, no prevalence setting and a restricted background. The background was filtered to include only the eastern rainforest corridor. X and Y axes represent the coordinates (WGS84). Black points represent the occurrence data. Future projections (2070) were estimated from three Global Circulation models (GCM) and two RCP scenarios. Top panels represent the most optimistic scenario (RCP26) and bottom panels represent the most pessimistic scenario (RCP85). Models produced similar results when setting prevalence to 0.5 (Equal total weights).

**Fig. 7.**
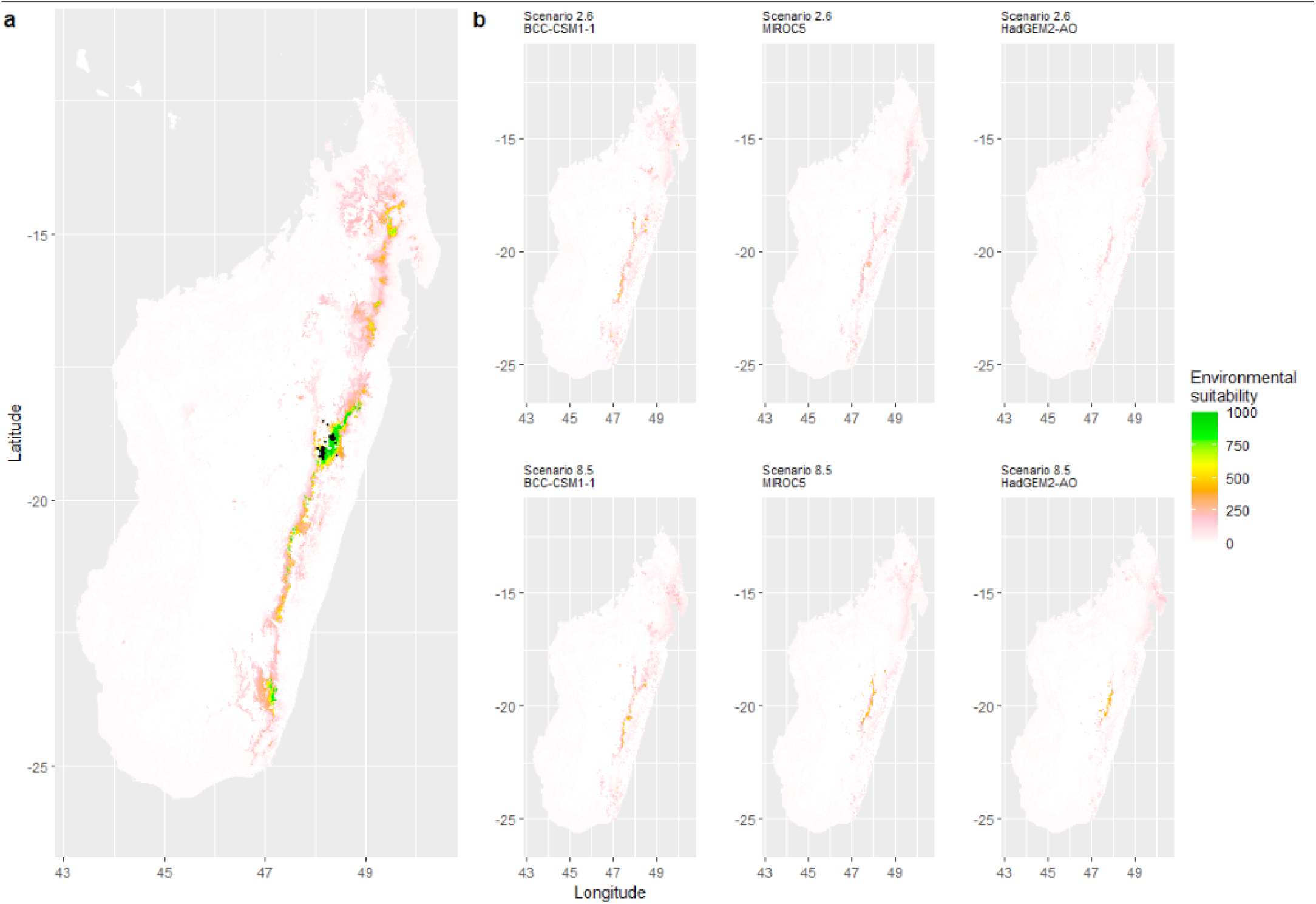
Current (a) and projected future (b) environmental suitability for *Mantella aurantiaca* using the Worldclim baseline climate. Predictions were filtered to include only forested areas. X and Y axes represent the coordinates (WGS84). Black points represent the occurrence data. Future projections (2070) were estimated from three Global Circulation models (GCM) and two RCP scenarios. Top panels represent the most optimistic scenario (RCP26) and bottom panels represent the most pessimistic scenario (RCP85).

**Fig. 8.**
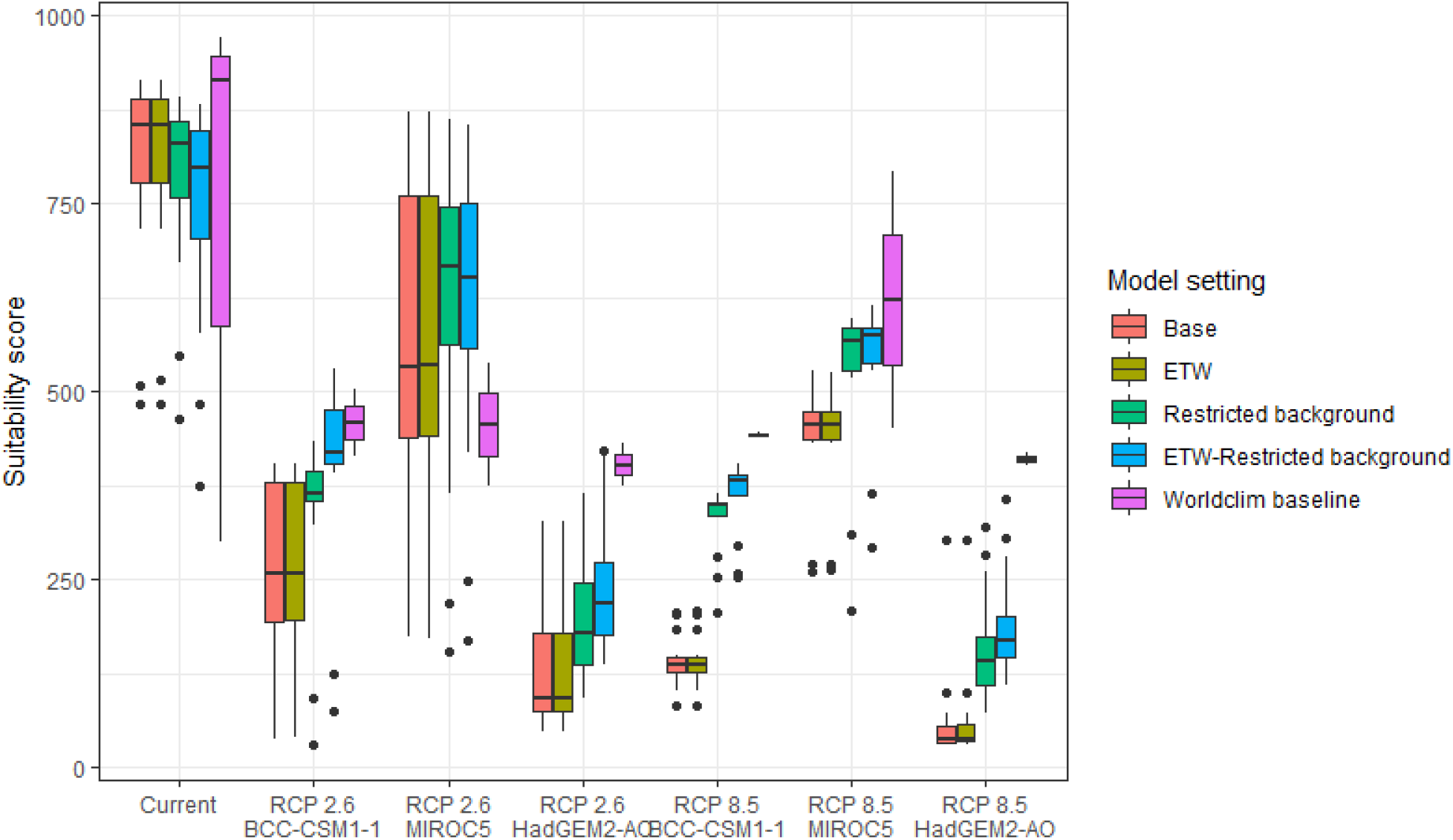
Variation in climate suitability scores at the occurrence points of *Phelsuma inexpectata* under current climate and 2070 climate. We show the variability in model predictions related to the climate scenario (RCP 2.6 versus RCP 8.5), the Global circulation model (3 GCMs), the background extent (wide versus restricted), prevalence setting (no setting versus Equal Total Weights, ETW) and baseline climate (Chelsa versus Worldclim). Each boxplot is composed of the first decile, the lower hinge, the median, the upper hinge and the nineth decile.

**Fig. 9.**
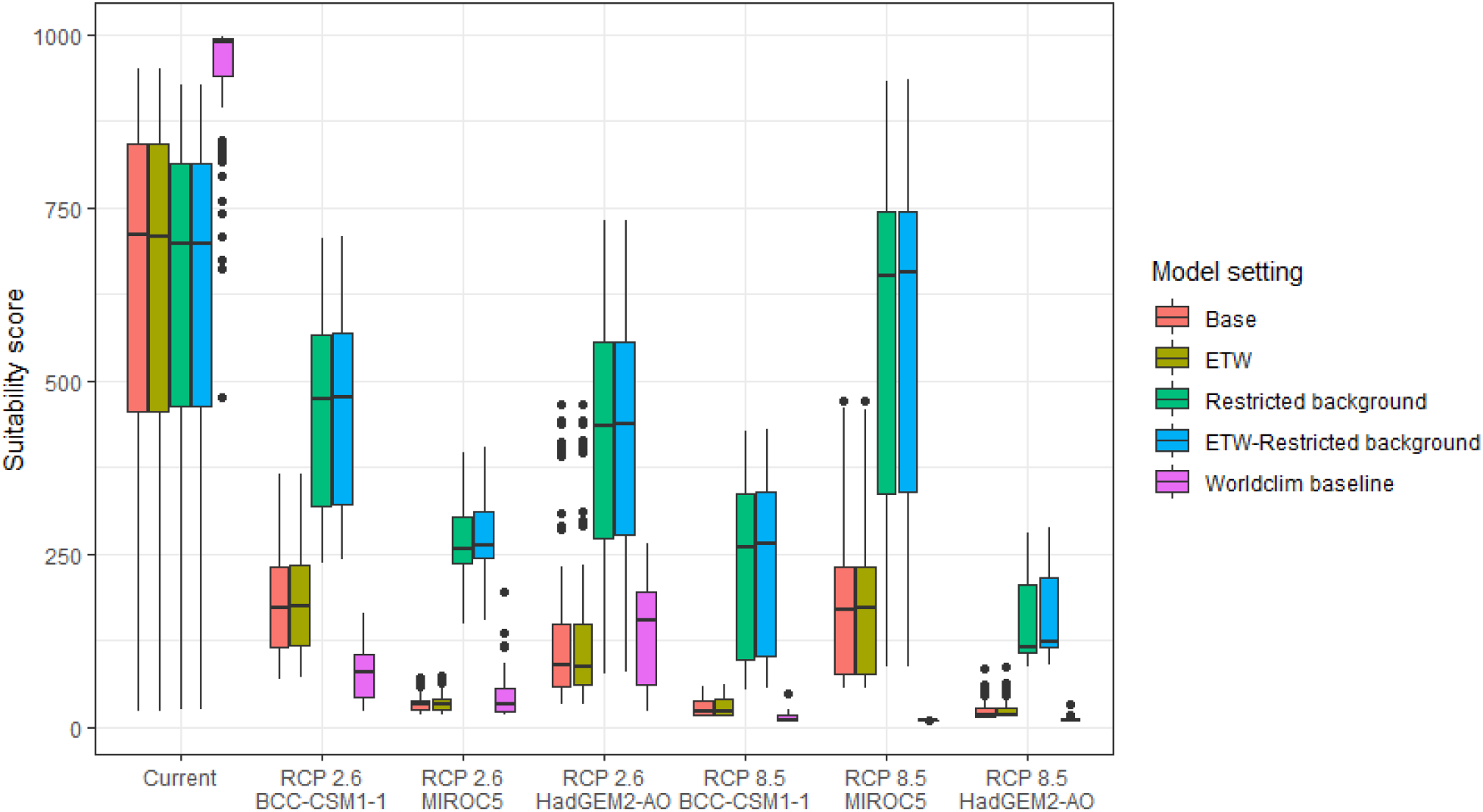
Variation in climate suitability scores at the occurrence points of *Mantella aurantiaca* under current climate and 2070 climate. We show the variability in model predictions related to the climate scenario (RCP 2.6 versus RCP 8.5), the Global circulation model (3 GCMs), the background extent (wide versus restricted), prevalence setting (no setting versus Equal Total Weights, ETW) and baseline climate (Chelsa versus Worldclim). Each boxplot is composed of the first decile, the lower hinge, the median, the upper hinge and the nineth decile.

In Reunion Island, predictions were the most variable between GCMs, but most ensemble models predicted a low climate suitability across the island. The highest suitability was found for the MIROC5 GCM, distributed along a band at higher altitudes (Fig. 2b). However, the MESS analyses indicated novel conditions in the most suitable areas, suggesting model extrapolations in these areas for this GCM (Fig. S13). According to the response curves (Fig. S14) and the predicted decrease in summer precipitations (Fig. S3c-d), climate suitability may have been overestimated.

In Madagascar, predictions varied the most with the background extent. Apart from the restricted background sets, none of the ensemble models identified any suitable area in the future. Nevertheless, we consistently identified a thin forest band in central Madagascar, south the current distribution, as the most suitable area across GCMs and scenarios (Fig. 3b). Models identified clear climatic windows, with no issue related to model extrapolation (Fig. S15, S16).

## Discussion

### Dramatic and widespread decline in climate suitability

We predict a strong decline of climate suitability in the whole current distribution area of both species by 2070. The subsequent high extinction risk in these species is not surprising, since small occupied areas are known to be good predictors of vulnerability to climate change (Pearson *et al*., 2014), but forecasting models were lacking and we contribute to fill this gap. We also predict that few or no zone will be suitable in the future across the entirety of their respective regions if those species are not given the opportunity to adapt. Depending on the Global Circulation model and the scenario, future projections still identified areas with suboptimal climatic conditions. In Reunion Island, the higher suitability found at ca. 20km west from the current distribution was presumably driven by an increase in precipitation. *Phelsuma inexpectata* is mostly present where precipitation is the lowest (< 500mm during the wet season, Fig., S10). The identified area coincides with the region with the driest conditions in Reunion Island in the future, according to these GCMs (Fig. S1). In all model sets, the MIROC5 GCM showed a higher—but suboptimal—climate suitability around the southern coast of the island, with the highest suitability along a band following the upper lands. This prediction is likely driven by the increase in overall temperature, along with reduced precipitation. The species lives under the hottest and driest conditions of the island, which fits the thermal requirements of the species for reproduction in captivity (28°C; McKeown, 1993). However, we believe this GCM is questionable because (1) the reduction in precipitation does not reflect the possible global increase in cyclone risk, and (2) the MESS analysis and response curves suggest an overestimation of the suitability scores. Note that this scenario (i.e. RCP 8.5)—which was the least pessimistic for *P. inexpectata—*is of interest to explore the widest range of possibilities in the future, but is considered unlikely (Hausfather & Peters, 2020).

In Madagascar, the current distribution is mainly explained by both winter (i.e. the dry season) and summer (i.e., the wet season) conditions with a short interval of suitability (Fig. S15). An important decline in climate suitability may therefore be driven by small changes in either winter or summer conditions. During winter, individuals migrate up the hills to shelter under dead woods and leaves and presumably enter a state of torpor or hibernation (Randrianavelona *et al*., 2010; Edmonds *et al*., 2020). The narrow climatic window may represent optimal conditions that minimise the risk of desiccation during this period of low activity. The species is also dependent on summer conditions, with a narrow window of suitable temperature and precipitations. This is highly consistent with a field study which recorded a surface temperature of 20-23°C at the occupied sites (Edwards et al., 2019) and an experimental design which found a decrease in activity when temperature deviates from 21.5°C (Edwards, 2019) with warm and rainy conditions (Fig. S12).

Summer corresponds to the period of reproduction, where females lay eggs under dead leaves on slopes, which are likely washed down into temporary pond during heavy rain episodes. This may explain the dependence with summer high precipitation and temperatures. This combination of climatic conditions is unlikely to be met in the future, except for the MIROC5 GCM under the RCP85 scenario (but this scenario is unlikely, Hausfather & Peters, 2020). However, we consistently identified potential suboptimal areas in the central-south of the eastern forest corridor and in one small area in the north east (corresponding to the top of mountains). For both species *P. inexpectata* and *M. aurantiaca*, the high level of habitat fragmentation associated to agricultural areas may limit potential distribution shifts, which calls the need for human intervention.

### Sources of uncertainty

The largest source of uncertainty was related to the GCM for *P inexpectata*, and to the background extend for *M. aurantiaca*. We followed a protocol that attempted to mitigate most sources of uncertainty, corresponding to the acceptable standards defined in Araújo *et al*. (2019). The long term, extensive and repeated efforts dedicated to species sampling enabled to define the current distributions of both species with high accuracy. Uncertainty map showed a high level of agreement between model replicates (i.e. modelling techniques, cross-validation runs and pseudo-absence runs; Fig. S17, S18).

In absence of information regarding the dispersal capacity of both species, we used two background corresponding to past dispersal hypotheses. The background covering the whole respective islands seems the most appropriate when assuming that only the sea could represent a dispersal barrier and that species could have dispersed throughout the whole island with regard to past climate and forest cover (Crottini et al. 2019), as assumed in Fieldsend et al. (2021). This might be the case for both species, especially *P. inexpectata* since the island is smaller and mostly occupied by a closely related species (*P. borbonica*; Dubos et al. 2021a). The restricted background corresponds to a hypothesis where *M. aurantiaca* dispersal is limited by current forest cover, and where *P inexpectata* is limited to the driest part of Reunion Island. This procedure enabled to encompass the widest range of possibilities with respect to two extreme hypotheses.

We included the 19 bioclimatic variables for the selection process, which is not recommended in most cases. However, we reduced the number of variables by removing the intercorrelated ones, and then selected the most biologically meaningful variables. We used the finest resolution available for climate data (i.e. 30 arc sec) which seems sufficient to discriminate suitable to unsuitable areas at the scale of Reunion Island and Madagascar. The inclusion of high-resolution land use variables enabled to improve the realism of both distributions and provides specific guidelines for conservation applications. In addition, we found a causal interpretation for the selected variables, which supports the biological significance of our models (Fourcade et al. 2018; Dubos et al. 2021b). An important limitation is that we did not use scenarios of future land use. This may lead to an overestimation of the available habitat in Madagascar, since the country is under important rates of deforestation (Veilledent et al., 2018). In Reunion Island, most of the natural habitats is incorporated in private properties or public gardens and in steep zones with limited access for agricultural practices. We believe that the apparent stability in agricultural areas would maintain the applicability of our results in the future. The inclusion of a range of different greenhouse-gas emission scenarios and GCMs showed an important uncertainty in the future suitability for *P. inexpectata*. However, this uncertainty is mostly related to one GCM (i.e. MIROC5), which we assume to be doubtful. The remaining ones consistently identified the most suitable area by 2070. Uncertainty related to model design was mitigated by the limitation of model complexity (with the ESM approach and the post-filtering technique), the removal of collinearity and the testing of a range of input parameters (number of pseudo-absences sets and crossvalidation subsets). Model performance was assessed with random partitions for *M. aurantiaca*, while spatial partitions are recommended. However, we argued that this methods is not appropriate for highly localised species due to strong imbalance between spatial blocks. We used multiple evaluation metrics, including discrimination (AUC and TSS) and reliability (i.e. calibration; Boyce index) metrics, all showing a high performance overall. We did not account for species dispersal ability, because the purpose was not to predict potential shifts, but to assess how suitable will be the climate in the future in order to identify candidate sites for restoration and translocation. Both species have low dispersal ability and their habitat is highly fragmented. Therefore, the potential for distributional shifts may be strongly limited and future projections must not be interpreted as the future distributions of our study species. An important limitation may be the absence of empirical knowledge on species thermal tolerances or other features of their climatic niche, adaptability and plasticity, which prevents us from determining whether our future projections underestimated the environmental suitability. Similarly, the extent of the species’ fundamental niche is unknown, and differs from the realised niche as a result of competition, habitat loss and environmental history, which may particularly important in endemic narrow-ranging species.

### A glimmer of hope

Despite the predicted low suitability in climate conditions, it is possible that species persist under changing conditions through adaptation or plasticity (Chevin *et al*., 2010; Hoffmann & Sgró, 2011). This hypothesis is supported by the persistence of two introduced populations of *P inexpectata* away from their current range in suboptimal environments (Fig. 2). Their persistence may result either from physiological or behavioural adaptation while benefiting from a combination of urban island effect and access to microclimate refuges in anthropogenic structures, as it is the case for *Phelsuma grandis* in Florida (Fieldsend *et al*., under review). It is also frequent that the current distribution of a given species represents only a fragment of its climatic niche (e.g., Guisan *et al*., 2014). For instance, experimental design on *M. aurantiaca* showed no expression of a thermal stress during periods of extreme heat (Edmonds *et al*., 2015). Therefore, predictions of climate suitability may underestimate the bounds of our model species niches. Further studies are needed to better characterise their climatic niche, and explore potential adaptive and plastic responses to changing conditions in these species and, more generally, in other threatened narrow-ranging species. Meanwhile, we encourage practitioners to implement conservation measures to grant those species a chance to adapt and persist. Nevertheless, the rate of climate change is generally faster than that of animal adaptive responses (Radchuk *et al*., 2019), which stresses the need for urgent actions. In the tropics, extinction risks may be greatly reduced with the development of land conservation programs provided climate change is mitigated by intergovernmental actions (Hannah et al., 2020).

### Conservation application

In Reunion Island, we identified two potential areas suitable for habitat restoration of *Phelsuma inexpectata* around the south-western coast and along a mid-altitude band. Despite climate conditions are predicted to be suboptimal, we believe these might represent the best options to ensure the long-term persistence of the species. Habitat restoration should be focussed on natural habitats invaded by non-native plants and in urbanised areas. Habitat restoration will consist in promoting the spread of native species (*Pandanus utilis, Latania lontaroides, Scaevola taccada* and *Psiadia retusa)* for which the species depends on (Bour, 1995). Finally, natural colonisation of newly suitable environments will be likely impossible for the species due to its poor dispersal ability and the high level of habitat fragmentation. We therefore encourage the design of translocation programs in management schemes.

In Madagascar, the area with the highest future climate suitability is located at the south of the current distribution, along the eastern rainforest corridor between the current distribution area and the north-west of the Vatovavy-Fitovinany region. The forest cover within the distribution range of *M. aurantiaca* has experienced a continued decline, regardless of the conservation status of the inhabited area (Piludu *et al*., 2015; Vieilledent *et al*., 2018). We recommend to reinforce the level of protection and to improve governance and management of the Mangabe (Moramanga region) and the Marolambo (Vatovavy-Fitovinany region) reserves. This can be achieved by promoting the development of alternative economic solutions through the development of valuable and sustainable activities that mitigate the rate of conversion of natural areas, the long-term management of soil fertility, and by considering the development of ecotourism. This species was included into a program that enabled to develop amphibian husbandry capacities in Madagascar, and that succeeded to establish captive bred colonies for this species (Rakotonanahary et al. 2017). Efforts of captive husbandry should be maintained (and possibly expanded to other species) and we encourage the design of translocation programs accounting for both local habitat characteristics and future climate suitability. Further study of biotic interaction between *M. aurantiaca* and other amphibians is needed to assess the consequences of its introduction.

### Are all endangered narrow-ranging species doomed to extinction?

Narrow-ranging species usually live under very specific environmental conditions and are the most vulnerable to climate change (Botts *et al*., 2013). Not only they are more prone to face distribution shifts and range contractions, but they might also disappear due to the strong alteration of their climatic envelope throughout their entire region. This may be the case for most narrow-ranging species with a specialised niche in regions with heterogeneous climates and high levels of endemism such as Madagascar but also Central America, South East Asia, tropical islands and more generally in tropical rainforests and mountains (Kier *et al*., 2009). In such regions, small shifts in climatic conditions may induce important changes in local environmental suitability for endemic—often specialised—species (e.g., Raxworthy *et al*., 2008). The risk is greater in tropical regions where species live closer to the upper bound of their thermal tolerance (Tewksbury *et al*., 2008; Şekercioĝlu *et al*., 2012; Dubos *et al*., 2019). Species that are already under threats are at the greatest risk because of a large array of synergistic effects (Şekercioĝlu *et al*., 2012). This might be the case for our two model species, which are threatened by habitat destruction and invasive species (Dubos, 2013; Piludu *et al*., 2015). Among the few Critically Endangered species for which the impact of climate has been investigated, most predict severe reductions in the climate suitability (e.g., Alamgir *et al*., 2015; Zhang *et al*., 2020), with the suitable range of the giant salamander *Andrias davidianus* predicted to decrease by more than two thirds by 2050 (Zhang *et al*., 2020). This species has faced a number of threats and climate change may be leading to imminent extinction despite a larger distribution range. Despite recent efforts developed for the monitoring of rare species in the tropics (e.g., Dubos et al., 2020), there is still an important lack of species occurrence data in these regions (Feeley & Silman, 2011). With the newly available high-resolution climate and land use data, the spectre of eligible species for SDMs has enlarged. We urge filling this data void by starting to assess the effect of climate change for narrow-ranging species at broader taxonomic scales, promote field investigations, assess species thermal requirements, and develop proactive conservation actions.

## Acknowledgements

We are grateful to all the field workers and their helpers who collected the data. We thank Boris Leroy for useful discussions and for his continued support.

## Data availability statement

The dataset and the scripts used in this analysis are available as R objects in the supporting information.

